# FGF activates Wnt signaling during newt lens regeneration

**DOI:** 10.64898/2026.11.20.768161

**Authors:** Leo Napoleon, Jacob Nipoti, Konstantinos Sousounis

## Abstract

Newts regenerate their eye lens following its complete removal. To accomplish this, pigmented epithelial cells located in the dorsal iris transdifferentiate to lens epithelial cells. FGF and Wnt signaling pathways have been implicated during this process, but their association remained unclear. Here, we used FGF signaling inhibitors to stall lens regeneration and studied the affected downstream mechanisms using immunofluorescence, western blotting, and RNAseq. We showed that entry to the cell cycle was slowed, ERK levels were lower, and several genes were differentially regulated with FGF inhibition. Among the impacted genes were Wnt signaling modulators revealing a potential connection between the pathways. FGF inhibition also stalled the gradual increase in beta-catenin abundance in the nucleus and junctions in iris pigmented epithelial cells during lens regeneration. To place Wnt downstream of the FGF signaling pathway, we pharmacologically activated Wnt and showed that it is sufficient to rescue the lens regeneration stalling phenotype observed with FGF inhibition. Overall, our data indicate that FGF activates the Wnt signaling pathway during newt lens regeneration.

## Introduction

Newts have the unique ability to faithfully regenerate the eye lens after complete surgical removal by transdifferentiating iris pigmented epithelial cells (IPECs) to lens epithelial cells (LECs)^1,2^. Fibroblast growth factor (FGF) signaling is known to have critical roles in eye and lens development^3,4^, and in tissue regeneration in a variety of model systems^5^. It is also a crucial part of the newt lens regeneration mechanism^6,7^. Notably, FGF2 injection in the intact newt eye triggers lens regeneration in the presence of the original lens^8^. In addition, mitotoxin-FGF2 conjugates and FGF receptor (FGFR) inhibition stall lens regeneration^7^. These suggest that iris cells expressing FGF receptors are necessary for the process and FGF signaling is one of the initiating steps. FGF2 is also necessary for lens regeneration in *in vitro* cultured IPECs^9–11^. Wnt signaling was implicated at later steps during the process with Wnt inhibitors inhibiting and Wnt ligands accelerating lens growths^11^. Overall, these studies strongly implicated FGF and Wnt signaling as necessary for lens regeneration, but little is known about their mechanistic relationship and their temporal function.

To address this gap, we first troubleshooted an experimental design using pharmaceuticals to perturbate FGF signaling and test the ability of iris cells to initiate lens regeneration following different durations of treatment. To gain a functional understanding of the role of FGF during newt lens regeneration, we assessed the cell cycle using immunofluorescence and downstream targets using western blotting, immunofluorescence, and RNA sequencing (RNAseq). The data indicated that the Wnt signaling was affected following FGF inhibition. To that end, we showed that activation of the Wnt signaling pathway can rescue the lens regeneration-stalling phenotype of FGF inhibition and restore certain IPEC functions. These findings provide for the first time a more direct connection between the FGF and Wnt signaling pathways during newt lens regeneration.

## Results

### FGF signaling is required early during lens regeneration

FGF inhibitors used in past studies showed variable outcomes^12^, thus, we first thought to identify small molecules and their effective concentration that could yield robust and predictable phenotypes. This is especially important since limiting phenotypic variability could improve mechanistic insights. Futibatinib, an irreversible pan FGFR inhibitor^13^, and erdafitinib, a reversible pan FGFR inhibitor^14^, were good candidates showing great tissue penetration and relatively low concentrations of action. To determine the optimal treatment concentration, newts were subjected to different amounts of erdafitinib following lens removal and found that lens regeneration was inhibited at 5uM but not at 0.5uM (Figure 1A). To gain information on the temporal window that FGF signaling is active during lens regeneration we continuously treated newts using the FGF inhibitors through 4 days post-lens removal (4dpl; Figure 1B and Figure 1C) and 10dpl (Figure 1D and Figure 1E). Following the treatment, newts were left to recover for at least 16 days, enough time for a lens vesicle to develop, before histology was performed to assess the phenotype. Newts treated with erdafitinib for 4 days developed smaller lens regenerates than DMSO controls but comparable to newts whose lenses were removed at day 4 (Figure 1C). This indicated that FGF signaling is engaged during the first 4 days following lens removal and can fully activate lens regeneration if the inhibition treatment is withdrawn. In contrast, newts treated with erdafitinib for 10 days developed considerably smaller lenses than newts whose lenses were removed at day 10 (Figure 1E). This indicates that newt iris cells were less potent in engaging the FGF signaling between 4 and 10dpl and were delayed or in a few cases unable to activate the lens regeneration process. Newts treated for up to 10dpl still reengaged regeneration, albeit delayed, suggesting that the eye maintains a pro-regenerative environment for prolonged periods of time post-lens removal (Figure 1E). Newts treated with the irreversible FGFR inhibitor futibatinib had an exacerbated phenotype with smaller lens regenerates than erdafitinib (Figure 1C and Figure 1E). The difference in regeneration speed may be attributed to the time required to turnaround FGFRs and reengage the FGF signaling. Overall, our data indicates that iris cells can initiate lens regeneration following inhibition of the FGF signaling pathway for at least 10dpl, however, regeneration rate is diminished with treatment of more than 4 days. These experiments provide additional insights into the temporal engagement and effectiveness of the FGF signaling during lens regeneration.

**Figure 1.**
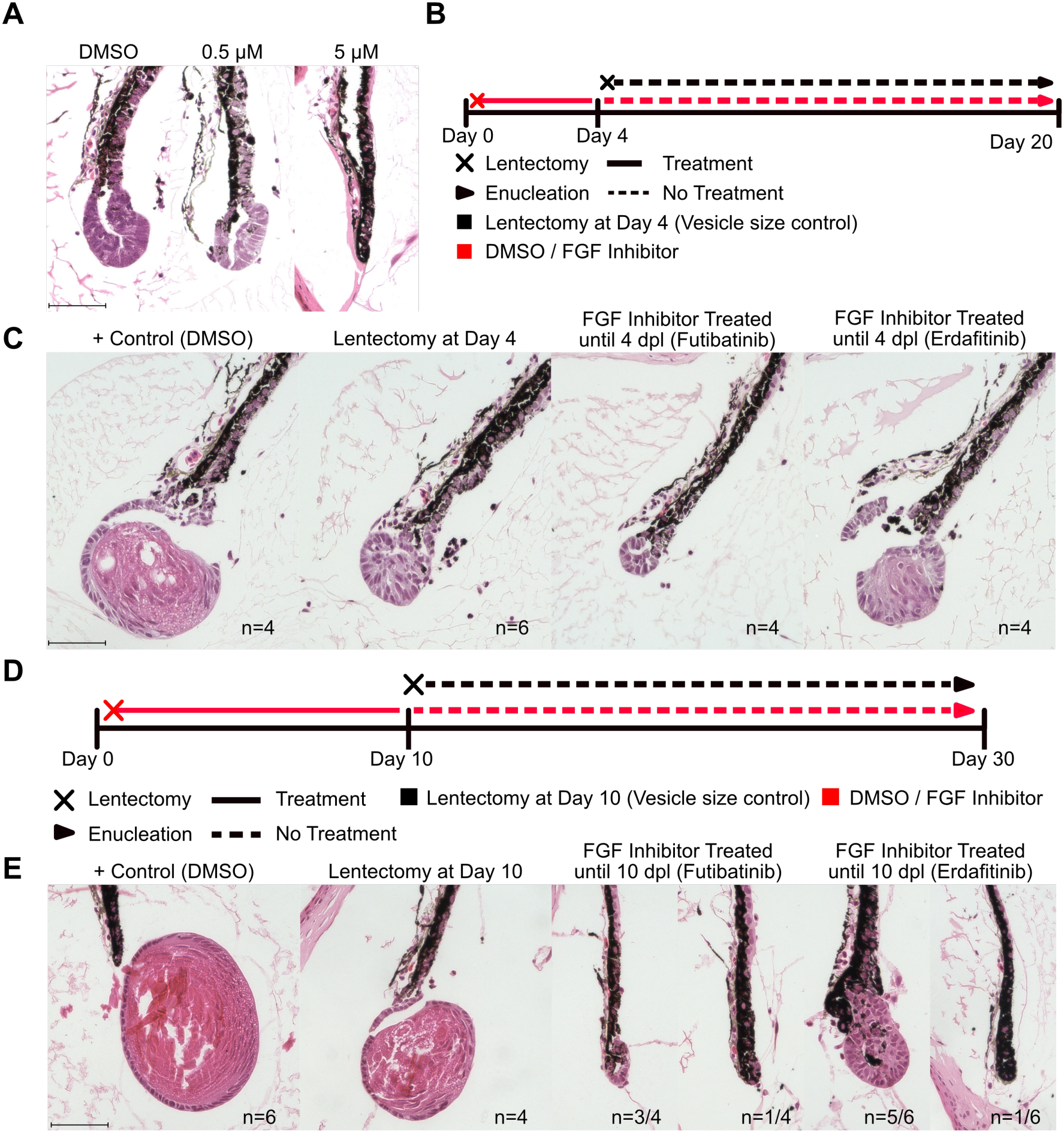
Lens regeneration is triggered even following 10 days of FGF inhibition but is most critical for the first 4 days post-lens removal. A) 14dpl dorsal iris sections from newts treated with DMSO and, 0.5 and 5 µM erdafitinib stained with Hematoxylin & Eosin (H&E). Newts treated with 5 µM erdafitinib lack lens vesicle formation. Scale bar 100 µm. B) Experimental design to test FGF inhibition up until 4dpl followed by 16 days of no treatment. A regeneration control group had their lenses removed at day 4 for comparison. C) Dorsal iris sections stained with H&E from B. Four days of FGF inhibition delayed but did not inhibit lens regeneration. Erdafitinib-treated newts developed comparable lens regenerates to the regeneration control group. As an irreversible pan FGFR inhibitor, Futibatinib inhibited lens regeneration more than erdafitinib. Scale bar 100 µm. D) Experimental design to test FGF inhibition up until 10dpl followed by 20 days of no treatment. A regeneration control group had their lenses removed at day 10 for comparison. Scale bar 100 µm. E) Dorsal iris sections stained with H&E from D. Lens regeneration was significantly delayed to initiate following withdrawal of the FGF inhibitors at 10dpl. Some eyes were not able to activate lens regeneration. Futibatinib yielded smaller lens vesicle than erdafitinib. Scale bar 100 µm.

### Pharmacological inhibition of FGF signaling stalls lens regeneration by reducing ERK and entry to the cell cycle

Newts treated with erdafitinib and futibatinib until 14dpl showed no signs of lens regeneration in all eyes tested (n=4-6, Figure 2A and Figure 2B). Erdafitinib was chosen for all downstream mechanistic experiments since futibatinib appeared to have lingering inhibiting effects for many days post treatment withdrawal (Figure 1). To confirm the predicted on-target mechanism of action of erdafitinib, we performed western blot analysis for total and phosphorylated p44/42 ERK, a known downstream effector of the FGF signaling^15,16^, between DMSO and erdafitinib treated iris tissue at 8dpl (Figure 2C). Erdafitinib treatment reduced total and phosphorylated ERK levels relative to alpha-tubulin in all four biological replicates, indicating that it is affecting FGF signaling in our system. A known function of the FGF-ERK pathway is cell cycle entry^17^, and IPECs in our erdafitinib-treated eyes appeared considerably smaller than those in DMSO-treated eyes (Figure 2B), suggesting a related phenotype in our model. To test this, we pulsed newts with the nucleotide analogue EdU four hours before collection at 7dpl and quantified EdU+ and phospho-Histone H3+ (pH3+) IPECs in histological sections (Figure 2A and Figure 2D). FGF-inhibited irises had significantly fewer EdU+ IPECs (t-test; p<0.01, Figure 2E) but similar amounts of pH3+ IPECs (t-test; p>0.05, Figure 2F). These data indicated that FGF signaling is required for IPEC cell cycle entry at the selected 7dpl timepoint. Overall, these data indicate that continued FGF inhibition completely stalled lens regeneration by dampening ERK signaling and blocking IPEC cell cycle entry.

**Figure 2.**
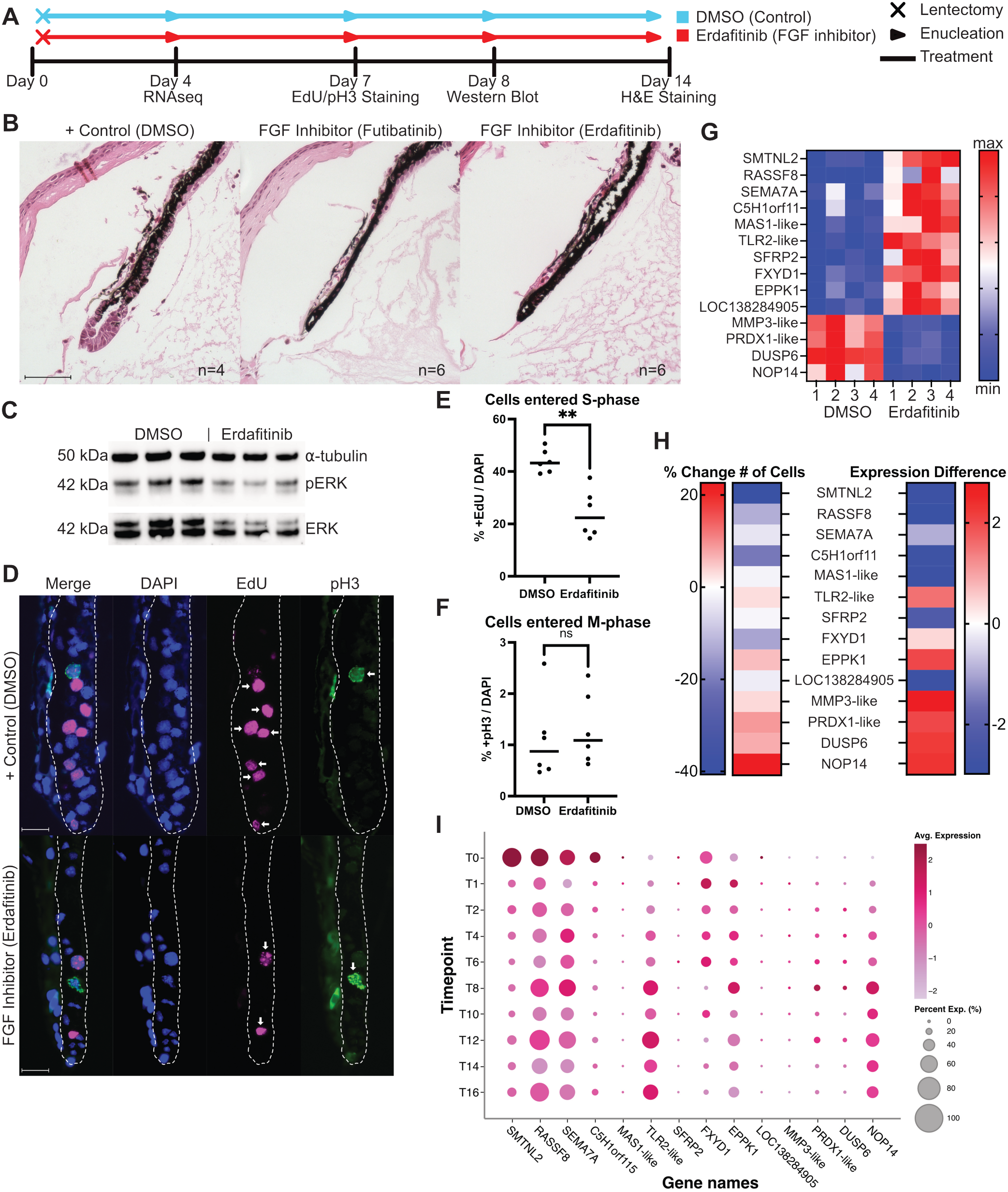
Inhibition of FGF signaling reduces ERK levels, slows cell cycle dynamics, and stalls the iris to an intact-like transcriptomic state. A) Experimental design of FGF inhibition and tissue collection days for downstream analysis. Tissues were collected at 4dpl for RNA extraction, 7dpl for EdU/pH3, 8dpl for protein, and 14dpl for histology. B) Dorsal iris sections stained with H&E at 14dpl. FGF inhibitors futibatinib and erdafitinib completely inhibited lens regeneration and iris enlargement. Scale bar 100 µm. C) Western blot for total and phosphorylated p44/p42 ERK, and α-tubulin under FGF inhibitor erdafitinib and DMSO treatments. Total and phosphorylated ERK levels were decreased under FGF inhibitor treatment. D) Representative images of dorsal iris sections at 7dpi stained with DAPI, EdU, and pH3. White dotted line indicates the iris pigmented epithelium which excludes the stroma in the anterior. Positive EdU/pH3 cells are marked with arrows. Scale bars 25 µm. E) Quantification of %EdU+/DAPI+ IPECs from D. There are less IPECs in the erdafitinib group entering S phase at 7dpl (Homoscedastic unpaired Student’s t-test, p<0.01). F) Quantification of %pH3+/DAPI+ IPECs from D. There is no change in IPECs in the erdafitinib group at the M phase at 7dpl (Homoscedastic unpaired Student’s t-test, p>0.05). G) Relative expression of transcripts found differentially regulated (q<0.05) in dorsal irises treated with erdafitinib and DMSO using RNAseq at 4dpl. FGF inhibited dorsal iris samples show regulation of FGF and Wnt signaling-related transcripts. H) Gene expression changes at 4dpl compared to intact iris derived from single cell RNA sequencing data during normal lens regeneration^18^. Percent change in number cells and expression difference indicated for genes identified in G. The expression difference indicates that FGF-inhibited irises display an intact-like transcriptomic profile. I) Dot plot showing expression of genes from G during lens regeneration. Data derived from a single cell RNA sequencing study^18^. ns p>0.05, *p<0.05, **p<0.01, ***p<0.001

### FGF-inhibited dorsal irises exhibit an intact-like transcriptomic profile and fail to activate the Wnt signaling

To better understand the role of FGF signaling during lens regeneration, we performed RNAseq on DMSO- and erdafitinib-treated dorsal iris tissue at 4dpl in four biological replicates. The use of an early timepoint could reveal the very first expression changes regulated by the FGF signaling and minimize other variables. Computational analysis revealed 14 genes to be differentially regulated (q<0.05) between treatments (Figure 2G). Among them, SMTNL2 was found significantly up-regulated in the FGF inhibited group. Interestingly, SMTNL2 was previously indicated as an intact iris gene marker in a single cell RNA sequencing (scRNAseq) study^18^. This suggested that the transcriptomic profile of the FGF-inhibited irises may be similar to that of intact iris. To test this, we used the timecourse expression data for 0 and 4 dpl from the scRNAseq study and assessed how the 14 identified genes changed over these timepoints during normal lens regeneration (Figure 2H). With the exception of FXYD1, EPPK1 and TLR2-like, all other 11 genes changed their level of expression in a pattern reflecting the early activation of lens regeneration. For example, expression of RASSF8 is reduced at 4dpl compared to intact during lens regeneration in both the number of cells expressing it and its overall levels (Figure 2H and Figure 2I). RASSF8 levels were found elevated in the FGF-inhibited irises at 4dpl (Figure 2G). Expression of DUSP6, a negative feedback inhibitor of MAPK and downstream target of FGF^19^, is increased at 4dpl compared to intact during lens regeneration (Figure 2H and Figure 2I). DUSP6 levels were found lower in the FGF-inhibited irises at 4dpl. These data indicated that FGF-inhibited irises show an intact-like transcriptomic profile. This observation is also corroborated at the histological level with FGF-inhibited IPECs failing to increase their cell size compared to controls (Figure 2B). Further analysis of the genes affected by the FGF inhibition treatment indicated a potential connection with the Wnt signaling pathway. For instance, SFRP2, a secreted inhibitor of Wnt^20^, is normally downregulated during lens regeneration (Figure 2H and Figure 2I). SFRP2 expression appeared significantly elevated under FGF inhibition potentially indicating that FGF signaling downregulates SFRP2 expression during lens regeneration to activate Wnt signaling (Figure 2G). Since Wnt signaling was previously shown to be important for newt lens regeneration^11^, we aimed to further explore the connection between FGF and Wnt.

### Wnt effector protein beta-catenin localizes at the nucleus and junctions during lens regeneration

To better understand the relationship between FGF and Wnt, we first thought to establish the temporal function of the Wnt signaling pathway during normal lens regeneration. To accomplish this, we performed immunofluorescence for beta-catenin, a well-characterized protein downstream of the Wnt signaling pathway^21–23^, in intact eye sections and during lens regeneration (Figure 3A). In the intact eye, beta-catenin was found to line the junctions of LECs in the intact lens (Intact lens; Figure 3A). Contrarily, IPECs had low abundance of beta-catenin in junctions and the nucleus (Intact iris; Figure 3A). After lens removal, beta-catenin progressively increased its abundance in the nuclei and junctions of IPECs (4 and 8dpl; Figure 3A). By 16dpl the beta-catenin staining pattern in the lens vesicle was similar to that of the intact lens (16dpl; Figure 3A). 16dpl IPECs entering the lens vesicle had increased beta catenin levels (16dpl, white arrows; Figure 3A). These data indicate that after lens removal beta-catenin gradually increases its presence in the nuclei and junctions of IPECs and eventually matches the staining pattern of the intact lens in the regenerated lens vesicle.

**Figure 3.**
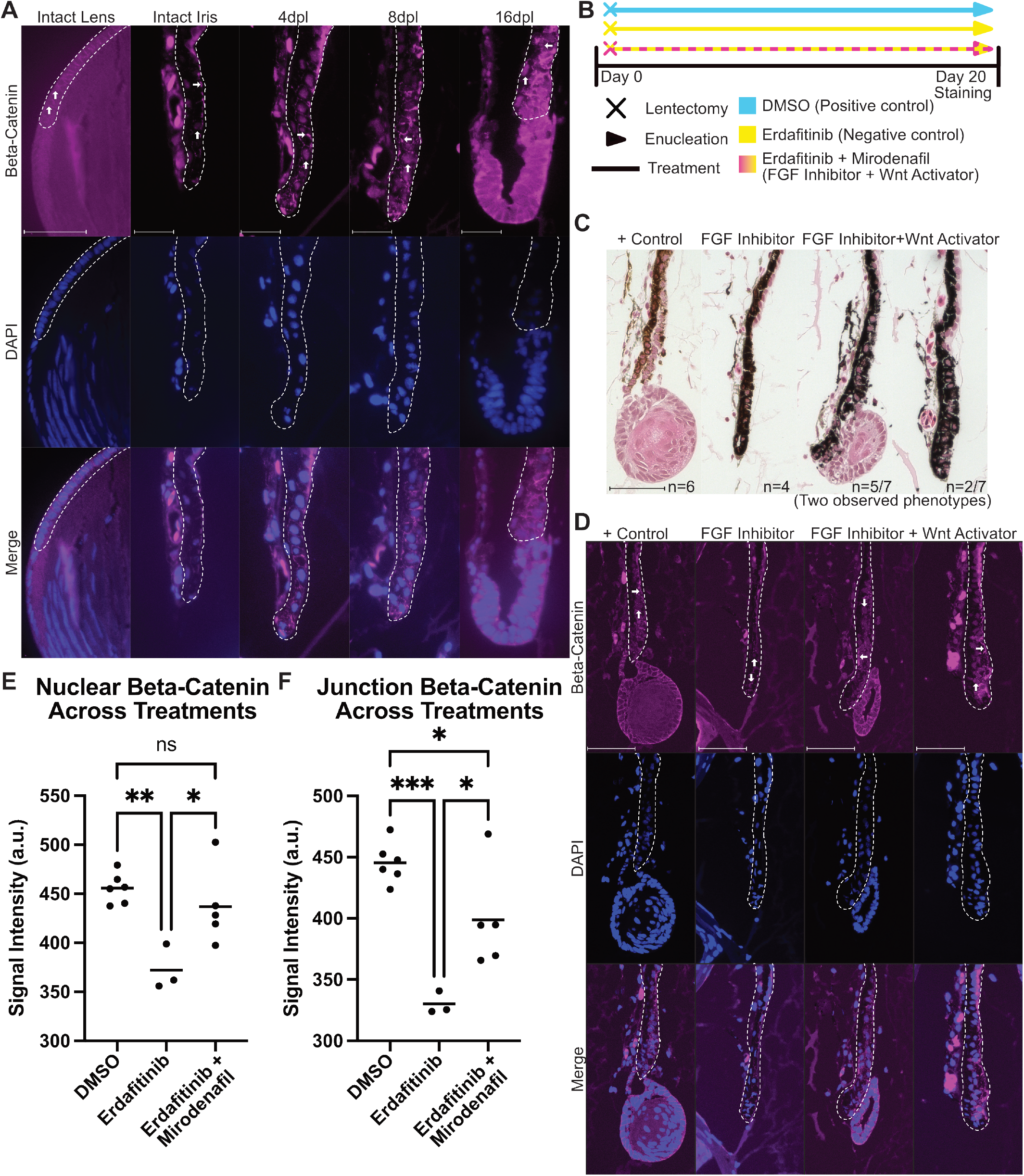
Wnt signaling is activated post-lens removal and can rescue the lens regeneration stalling phenotype under FGF inhibition. A) Beta-catenin immunofluorescence in the intact lens, intact iris and 4, 8, 16dpl irises. Arrows indicate beta-catenin localization in nuclei and junctions. Scale bar for intact lens 100 µm, scale bars for iris tissues 50 µm. B) Experimental design for FGF inhibition and Wnt activation. All samples were treated for 20dpl. C) Dorsal iris sections stained for H&E from B. DMSO treated eyes regenerated a lens (6/6). Erdafitinib treated eyes did not regenerate a lens (0/4). Erdafitinib and mirodenafil treated eyes regenerated a lens in 5/7 eyes. Scale bar 100 µm. D) Representative images of dorsal iris sections at 20dpl stained for beta-catenin and DAPI. The iris pigmented epithelium is circled, excluding the stroma. Arrows indicate beta-catenin in the junctions and nucleus. Scale bars 50 µm. E) Quantification of nuclear beta-catenin abundance in IPECs from D. Wnt activation under FGF inhibition resulted in higher beta-catenin levels in the nucleus than FGF inhibition alone (p<0.05), and no difference than DMSO (p>0.05) determined by one-way ANOVA with Tukey’s post-hoc multiple comparisons test. F) Quantification of beta-catenin abundance in IPEC junctions from D. Wnt activation under FGF inhibition resulted in higher beta-catenin levels in the junctions than FGF inhibition alone (p<0.05), but less than DMSO (p<0.001) determined by one-way ANOVA with Tukey’s post-hoc multiple comparisons test. ns p>0.05, *p<0.05, **p<0.01, ***p<0.001

**Figure 4.**
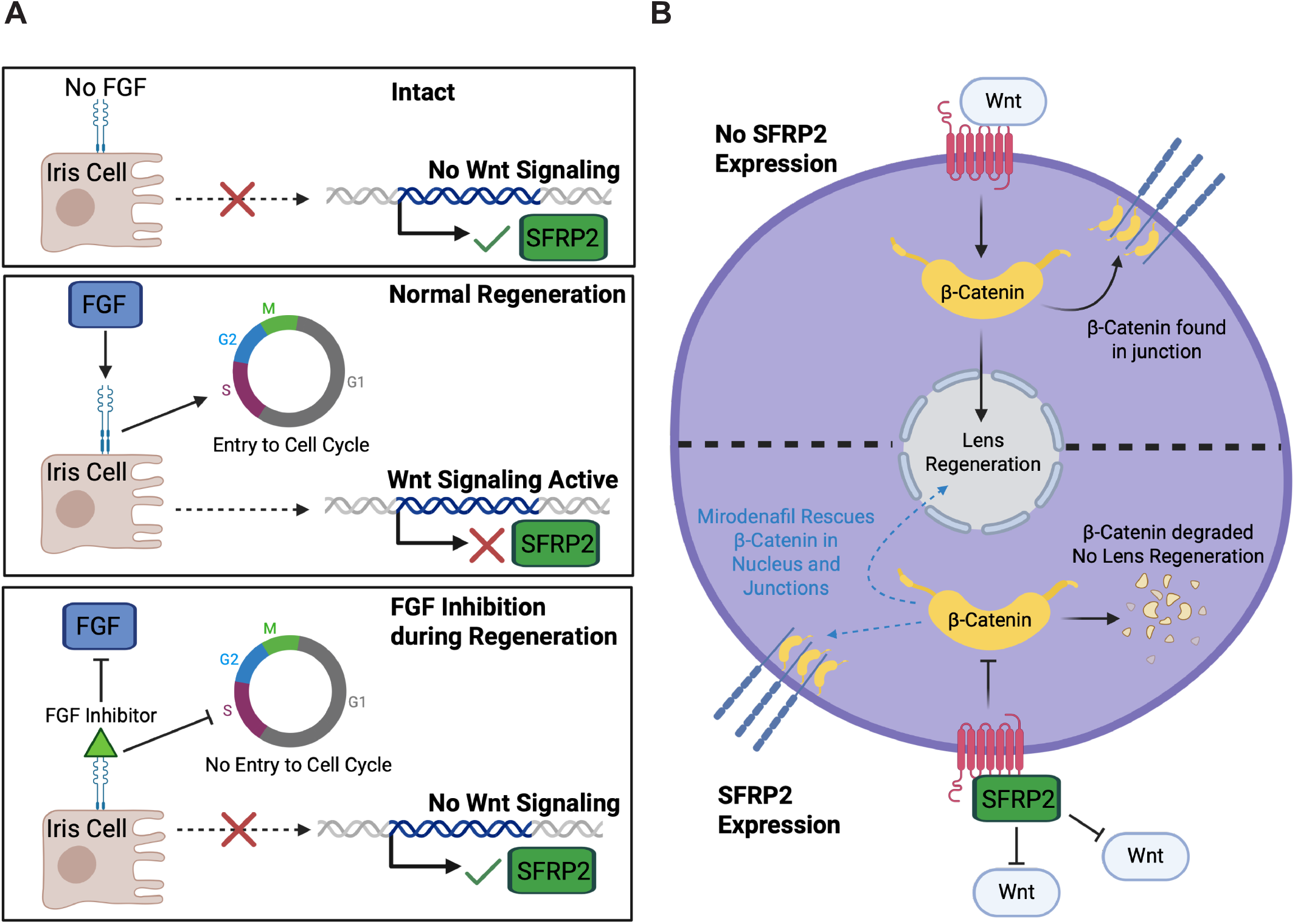
Cartoon schematic of the association between FGF and Wnt signaling pathways, and their role during lens regeneration. Created with BioRender. A) FGF signaling has roles in cell cycle entry and regulation of the Wnt signaling, for example, via SRFP2 expression during lens regeneration. Intact iris tissue has no FGF signaling and shows high SFRP2 expression. Upon lens removal, iris cells engage the FGF signaling which activates entry to the cell cycle and suppresses expression of the Wnt inhibitor SFRP2. Pharmacological inhibition of FGF signaling during lens regeneration slows cell cycle dynamics and keeps SFRP2 levels elevated. As a Wnt signaling inhibitor, regulation of SFRP2 downstream of FGF indicates, a previously unknown, link between the pathways. B) SFRP2 levels are elevated in intact and FGF-inhibited iris tissue which suppresses Wnt signaling and leads to beta-catenin degradation and no lens regeneration. During lens regeneration, Wnt signaling is engaged and leads to the localization of beta-catenin in the nucleus and junctions. Mirodenafil treatment activates the Wnt signaling and rescues beta-catenin localization and lens regeneration in the FGF-inhibited iris tissue.

### Pharmacological activation of Wnt signaling rescues lens regeneration in FGF-inhibited irises

We previously showed that FGF signaling inhibition using erdafitinib completely stalls lens regeneration in all animals tested (Figure 2B). Using RNAseq, we found that the FGF-inhibited dorsal iris failed to activate Wnt as indicated by the elevated expression of the Wnt inhibitor SFRP2 (Figure 2G). To test whether Wnt is downstream of the FGF signaling pathway, we used mirodenafil, a drug known to activate the Wnt signaling^24^, while simultaneously inhibited FGF signaling using erdafitinib until 20dpl (Figure 3B). As expected, all DMSO-treated, positive control, animals regenerated a sizable lens vesicle (DMSO, n=6 eyes; Figure 3C) and all erdafitinib-treated, negative control, animals failed to initiate regeneration in all samples (Erdafitinib, n=4 eyes; Figure 3C). Interestingly, mirodenafil treatment was able to rescue the erdafitinib-induced lens regeneration stalling phenotype in ~70% of the cases (Mirodenafil + Erdafitinib, n=5/7 eyes; Figure 3C). In the two eyes that regeneration was not rescued, there is a notable thickening of the iris versus the erdafitinib-only treated samples (Mirodenafil + Erdafitinib, n=2/7 eyes; Figure 3C). These data strongly indicate that Wnt signaling is downstream of FGF signaling. To better understand the mechanism of action for our pharmaceutical treatments as it relates to the Wnt signaling, we stained the sections for the Wnt effector protein beta-catenin (Figure 3D). Quantification of beta-catenin abundance in IPECs showed that erdafitinib treatment significantly lowered the amounts of beta-catenin in the nucleus (p<0.01; DMSO vs erdafitinib, p<0.05; erdafitinib vs mirodenafil/erdafitinib; Figure 3E) and junctions compared to DMSO and mirodenafil treatments (p<0.001; DMSO vs erdafitinib, p<0.05; DMSO vs mirodenafil/erdafitinib; Figure 3F). Mirodenafil treatment rescued the beta-catenin levels in the nucleus (p>0.05; DMSO vs mirodenafil/erdafitinib; Figure 3E) and partially rescued the levels in the junctions (p<0.05; DMSO vs mirodenafil/erdafitinib; Figure 3F) compared to erdafitinib-only treatment. These data indicate that erdafitinib treatment inhibits the activation of beta catenin-mediated Wnt signaling while mirodenafil treatment activated Wnt signaling in IPECs. Mirodenafil treatment was also sufficient to rescue the lens regeneration stalling phenotype observed with the FGF-inhibiting erdafitinib treatment by restoring beta catenin levels in IPEC nucleus and junctions. Overall, we provide strong evidence that FGF signaling activates the Wnt signaling pathway during newt lens regeneration.

## Discussion

Here we describe for the first time that Wnt signaling is activated by the FGF signaling during newt lens regeneration. To establish this connection, we first troubleshooted a highly reproducible lens regeneration stalling phenotype *in vivo* with two FGFR inhibitor pharmaceuticals, erdafitinib and futibatinib (Figure 2B). This was a critical foundation lacking from past studies whose regiments used were yielding phenotypes ranging from no regeneration to visible lens vesicle formation, thus complicating analysis of downstream experiments. By completely inhibiting lens regeneration in all eyes tested, we were able to better understand the role of the FGF signaling pathway. In line with past studies, we show that FGF inhibition stalls cell cycle entry at 7dpl (Figure 2D), a timepoint known to include cycling IPECs^18,25,26^. Using RNAseq in 4dpl irises, we aimed to capture the early effects of FGF signaling in the transcriptomic landscape. This analysis revealed just 14 differentially regulated genes and given the early timepoint sampled and the striking phenotype of the inhibitor, these genes likely reflect direct or secondary targets of the FGF signaling pathway. One of the genes, DUSP6, a negative feedback regulator of FGF, is, in fact, known to be directly downstream of FGF signaling^19^. DUSP6 is known to be regulated by ERK, a FGF effector protein found to be decreased with our FGF inhibitor treatment (Figure 2C). The most interesting genes, however, were those associated with the Wnt signaling pathway. In particular, SFRP2, a secreted Wnt inhibitor^20^, stayed elevated with FGF inhibition indicating that FGF signaling may act to disinhibit the Wnt signaling during lens regeneration (Figure 2G). Past studies using *in vitro* newt lens regeneration models had shown the importance of FGF and Wnt during the process and had proposed a two-step mechanism with FGF early and Wnt later^11^. Most notably, Wnt activation could induce lens regeneration from the incompetent ventral iris while Wnt inhibition could stall regeneration from the dorsal iris. However, similar experiments *in vivo* were unsuccessful and rescue experiments were not performed. To that end, and using our very reproducible FGF inhibition treatment phenotypes as a foundation, we activated the Wnt signaling pathway using mirodenafil *in vivo*^*24*^. This treatment rescued the lens regeneration stalling phenotype caused by the FGF inhibition, strongly indicating that Wnt is downstream of FGF in this model system. Mirodenafil treatment also restored the beta-catenin localization pattern in the iris which further supports the on-target mechanism of this drug. Overall, our inhibition and rescue experimental design provides for the first time a platform to further study the intricate molecular mechanisms that enable IPECs to reprogram to LECs during newt lens regeneration.

Our study strongly supports past studies indicating FGF and Wnt as two signaling pathways that mark important early and late functions during newt lens regeneration^6–8,11,12^. By inhibiting FGF for 4 and 10dpl, and withdrawing the treatment, we were able to estimate the window of effectiveness of this signaling pathway. With 4 days of treatment, lens regeneration could fully restart the process while with 10 days of treatment, lens regeneration was significantly impacted. This indicates that IPECs need to engage FGF signaling early post-lens removal to maintain their optimal regeneration rate. Our RNAseq analysis also revealed injury-response genes including a matrix metalloproteinase, that was significantly downregulated, and a toll-like receptor, that was significantly elevated upon FGF inhibition. These temporally coincide with the highly inflammatory environment in the eye during the early phases of regeneration which may affect the effectiveness of the FGF signaling^18,27,28^. In addition, our RNAseq analysis strongly indicates that FGF inhibition keeps the iris in an intact-like state (Figure 2G-I). Eleven out of the fourteen differentially regulated genes following FGF inhibition showed expression levels matching that of intact iris. For instance, SFRP2 was elevated in the intact iris compared to post-lens removal, and appeared elevated with FGF inhibition treatment in-line with this hypothesis. The beta-catenin immunofluorescence staining pattern in FGF-inhibited irises was also similar to that of an intact iris (Figure 3D), further indicating that regeneration was completely stopped with these treatments. These data, together with past studies showing that a single injection of FGF2 can trigger lens regeneration in the presence of the original lens^8^, strongly position the FGF signaling pathway as the very first step that initiates newt lens regeneration. Mirodenafil treatment restored the beta-catenin staining pattern (Figure 3D) and rescued lens regeneration in FGF-inhibited irises (Figure 3C), indicating that Wnt signaling is sufficient to advance newt lens regeneration under these conditions. However, the location of the rescued regenerated lens vesicle is considerably different from that of DMSO controls. The rescued lens vesicle appeared towards the base and not at the tip of the iris which indicates that not all processes were rescued by Wnt activation (Figure 3C). One possibility is that early FGF activation is required for IPECs to enter the cell cycle to produce and position more cells at the tip of the iris for lens reprogramming. A different possibility is that the iris is still producing the Wnt inhibitor SFRP2 that targets the iris tip versus a more proximal location where the Wnt activator drug could better engage IPECs. Future studies will aim to further decipher the molecular mechanisms underlying this process, including genetic regulation of SFRP2 and the role of beta-catenin, that allow FGF and Wnt to interact and perform their roles during lens regeneration.

In summary, our study provides the first *in vivo* rescue experiment that strongly links the FGF and Wnt signaling pathways during newt lens regeneration. These data, along with our downstream functional analysis, pave the road for a deeper molecular and genetic understanding of how IPECs are activated and reprogrammed to LECs in this model system. This would further our understanding on how highly regenerative organisms can naturally regenerate lost organs and tissues.

## Materials and Methods

### Animal handling

Experiments involving animals were performed in accordance with the University of New Hampshire (UNH) Institutional Animal Care & Use Committee Research (IACUC) protocols #220604 and #241003. Animals were raised in dechlorinated tap water (housing water) at room temperature. All surgical procedures were carried out while animals were anesthetized with buffered 0.1% tricaine.

### Lens removal

Following confirmation of anesthesia, the cornea was punctured using a scalpel. Using surgical scissors, a cut across the length of the cornea was made. Fine point forceps were then inserted into the eye and used to gently pinch and remove the lens. Both eyes had their lenses removed. Animals were then placed directly into housing water pretreated with either experimental or control reagents.

### Iris removal

Following confirming of anesthesia, the side of the sclera just below the cornea/iris plane was punctured. Surgical scissors were used to cut out the anterior segment of the eye. Tissues were moved to cold 0.7X phosphate buffered saline without magnesium and calcium (0.7X PBS). The iris ring was then cleaned of cornea and adjacent tissues. Dorsal iris segments from two newts, four eyes, were used for the RNAseq samples. The entire iris ring from two newts, four eyes, were used for the western blot samples.

### Enucleation

At selected timepoints following lens removal and following daily pharmacological treatments, animals were anesthetized, and eyes were removed with the dorsal eyelid attached to determine proper orientation during embedding. The eyes were placed directly into 4% paraformaldehyde in 0.7X PBS overnight before tissue processing and embedding.

### Drug preparation and treatment

Final concentrations for each drug in treated tap water were 5 µM for erdafitinib (HY-18708; MedChemExpress), 1 µM for futibatinib (S8848; Selleckchem), and 10 µM for mirodenafil (HY-14930A; MedChemExpress). A 1000x stock concentration of each drug was made in dimethyl sulfoxide (DMSO) and diluted 1/1000 in housing water. DMSO was diluted 1/1000 for control groups. Housing water and treatments were replenished daily until the end of the experiment.

### Tissue processing, embedding, sectioning, and staining

After enucleation, eyes were dehydrated in an ethanol gradient to 100% followed by three xylene and paraffin washes. The eyes were then placed into a mold for sagittal sectioning. Sectioning was performed at 5 µm using a rotary microtome. For morphological analysis, ribbons of sections were placed onto slides and stained with hematoxylin and eosin to visualize the structure. Images were taken using an EVOS FL Auto 2 microscope.

### Immunofluorescence (IF), EdU, and DAPI staining on Formalin Fixed Paraffin Embedded (FFPE) Slides

After seven days of experimental treatments, newts were injected with EdU following established protocols^29^ and eyes were collected and fixed in 10% normal buffered formalin after a four-hour pulse. Eyes were processed as previously described. The FFPE slides were rehydrated and boiled in 1X citrate buffer for ten minutes followed by three 0.7X PBS washes. Blocking was performed with 2% Bovine Serum Albumin (BSA) in 0.1% Triton-X-100 for one hour followed by two washes of 0.7X PBS. Phosphohistone H3 antibody was added overnight in 2% BSA in 0.1% Triton-X-100 at 4ºC. Control slides did not receive primary antibody and instead were covered with 2% BSA 0.1% Triton-X-100 0.7X PBS. The slides were then rinsed with 0.7X PBS three times and secondary antibody in 2% BSA in 0.1% Triton-X-100 was added for 2 hours at room temperature. Slides were rinsed with 0.7X PBS three times before stained for EdU following the manufacturer’s recommendations. Slides were rinsed with 0.7X PBS three times before adding 1X DAPI in 0.7X PBS for 5 minutes at room temperature. The slides were rinsed with 0.7X PBS three times before mounted using Diamond ProLong Antifade and allowed to dry overnight at 4ºC. Slides stained for beta-catenin were prepared with Alexa Fluor™ 647 Dye secondary antibody and not stained for EdU. Double blind quantification of %EdU+/DAPI+ and %pH3+/DAPI+ was performed by counting cells within the iris pigmented epithelium. Antibodies used and their dilutions were beta-catenin (#8480, Cell Signaling Technology) 1:200, and Anti-phospho-Histone H3 Ser10 (06-570, Millipore Sigma) 1:400. Images were taken using a Nikon Eclipse Ni-E microscope.

### Quantification of beta-catenin in the nucleus and junctions from IF images

Fiji^30^ was used to manually indicate the region of interest (ROI) which contained only the iris pigmented epithelium and excluded the stroma, lens, and ciliary. Cellpose^31^ was used to detect nuclei based on DAPI fluorescence and measured signal intensity of the beta-catenin staining within the nucleus. Non-nuclear signal intensity within the ROI was used to estimate beta catenin in the junctions. Nuclear selections by cellpose in each image was manually examined to ensure accuracy before the data was used.

### Protein extraction, BCA assay, and Western Blot

After eight days of experimental treatments, extracted irises were dissolved in RIPA buffer with 1X phosphatase/protease inhibitors. BCA assay was performed to determine sample concentration and ensure equal loading of 7.54 µg per lane/ biological replicate. Gel electrophoresis was performed using a NuPage 4-12% Bis-Tris gel and gel transfer using a Bio-Rad trans blot turbo transfer using a nitrocellulose membrane. Blocking was performed with 5% BSA 0.1% Tween20 1X TBS (TBST) for one hour at room temperature. Primary antibodies were diluted in 5% BSA in TBST and incubated at 4ºC overnight. HRP linked secondary antibodies were diluted in 5% BSA in TBST and incubated for one hour at room temperature. The blots were washed three times for ten minutes with TBST after each incubation. ECL was used for five minutes before imaging using Biorad chemidoc imager. Antibodies used and their dilutions were phospho-ERK (#4370, Cell Signaling Technology) 1:2000, ERK (#9102, Cell Signaling Technology) 1:1000, alpha-Tubulin (#3873, Cell Signaling Technology) 1:1000, and anti-rabbit HRP (#7074, Cell Signaling Technology) 1:2000, and anti-mouse HRP (#7076, Cell Signaling Technology) 1:2000.

### RNA sequencing: Extraction and analysis

RNA from the isolated dorsal iris tissue was extracted using TRIzol reagent (Thermo Fisher Scientific) and RNA Concentrator (Zymo Research). RNA was processed at the University of New Hampshire Hubbard Center for Genome Studies with Kapa HyperPrep RNA and run on a NovaSeq SP PE 250 lane following sample QC. Reads were assessed with FastQC and trimmed using Trimmomatic-0.39^32^ with CROP:65 HEADCROP:12 LEADING:3 TRAILING:3 MINLEN:53 arguments. Differential gene expression analysis was performed using kallisto^33^ and sleuth^34^ packages against the *Pleurodeles waltl* RefSeq transcriptome^35^. Differential gene expression analysis included in Supplementary Table 1. Trailmaker® (Parse Bioscience) was used to generate Figure 3H and 3I with timecourse expression data from prior studies^18^ for the differentially regulated genes identified in the RNA sequencing experiment.

### Statistics

Homoscedastic unpaired t-test was used for data in Figure 2E and 2F. One-way ANOVA with normal distribution and Tukey post-hoc multiple comparisons test was used for data in Figure 3E and 3F.

## Acknowledgements

We would like to thank the staff of the University of New Hampshire’s (UNH’s) Animal Research Office, Dr. Dean Elder, Dr. Linnea Morley, and Meadow Miles for excellent animal care and assistance. We would also like to thank Olivia Williams for helpful discussions and technical assistance, and Dr. Nate Oldenhuis for assistance with the Biorad chemidoc imager. The EVOS and Nikon microscopes used in this study were purchased by the UNH’s Center for Integrated Biomedical and Bioengineering Research (CIBBR) through a grant from the National Institute of General Medical Sciences (NIGMS) of the National Institutes of Health (NIH) (Grant P20GM113131) and are managed by the UNH University Instrumentation Center (UIC). Research reported in this publication was supported by the National Eye Institute of the NIH under Award Number R00EY029361 and the NIGMS of the NIH under Award Numbers P20GM113131 to KS. The content is solely the responsibility of the authors and does not necessarily represent the official views of the NIH. Research was also supported by UNH’s Research Experience and Apprenticeship Program (REAP) and Summer Undergraduate Research Fellowship (SURF) to LN.

## Footnotes

### Author contributions

Conceptualization, supervision, and funding acquisition done by K.S.; Animal handling done by L.N., J.N., and K.S.; Surgeries done by J.N. and K.S.; Tissue processing, histological sectioning, staining, imaging, and analysis completed by L.N. and J.N.; Bioinformatic analysis completed by K.S. Visuals generated by L.N.; Text written and edited by L.N., J.N., and K.S.

### Data and resource availability

Raw and processed data will be available upon publication.

### Competing interests

The authors declare no competing interests.

## Table legends

Supplementary Table 1

Differential gene expression analysis for DMSO- and Erdafitinib-treated dorsal iris samples.

## Notes

### Competing Interest Statement

The authors have declared no competing interest.

## References

1. Reyer, R. W. Regeneration of the Lens in the Amphibian Eye. Q. Rev. Biol. 29, 1–46 (1954).

2. Tsonis, P. A. How to build and rebuild a lens. J. Anat. 209, 433–437 (2006).

3. Robinson, M. L. An essential role for FGF receptor signaling in lens development. Semin. Cell Dev. Biol. 17, 726–740 (2006).

4. McAVOY, J. W., Chamberlain, C. G., Iongh, R. U. De, Richardson, N. A. & Lovicu, F. J. The Role of Fibroblast Growth Factor in Eye Lens Development a. Ann. N. Y. Acad. Sci. 638, 256–274 (1991).

5. Farooq, M., Khan, A. W., Kim, M. S. & Choi, S. The Role of Fibroblast Growth Factor (FGF) Signaling in Tissue Repair and Regeneration. Cells 10, 3242 (2021).

6. Del Rio-Tsonis, K., Jung, J. C.Chiu, I.-M. & Tsonis, P. A. Conservation of fibroblast growth factor function in lens regeneration. Proceedings of the National Academy of Sciences 94, 13701–13706 (1997).

7. McDevitt, D. S., Brahma, S. K., Courtois, Y. & Jeanny, J.-C. Fibroblast growth factor receptors and regeneration of the eye lens. Developmental Dynamics 208, 220–226 (1997).

8. Hayashi, T., Mizuno, N., Ueda, Y., Okamoto, M. & Kondoh, H. FGF2 triggers iris-derived lens regeneration in newt eye. Mech. Dev. 121, 519–526 (2004).

9. Bhavsar, R. B., Nakamura, K. & Tsonis, P. A. A System for Culturing Iris Pigment Epithelial Cells to Study Lens Regeneration in Newt. Journal of Visualized Experiments https://doi.org/10.3791/2713 (2011) doi:10.3791/2713.

10. Hoffmann, A., Nakamura, K. & Tsonis, P. A. Intrinsic Lens Forming Potential of Mouse Lens Epithelial versus Newt Iris Pigment Epithelial Cells in Three-Dimensional Culture. Tissue Eng. Part C Methods 20, 91–103 (2014).

11. Hayashi, T., Mizuno, N. & Kondoh, H. Determinative roles of FGF and Wnt signals in iris-derived lens regeneration in newt eye. Dev. Growth Differ. 50, 279–287 (2008).

12. Del Rio-Tsonis, K., Trombley, M. T., McMahon, G. & Tsonis, P. A. Regulation of lens regeneration by fibroblast growth factor receptor 1. Developmental Dynamics 213, 140–146 (1998).

13. Ito, S. et al. Discovery of Futibatinib: The First Covalent FGFR Kinase Inhibitor in Clinical Use. ACS Med. Chem. Lett. 14, 396–404 (2023).

14. Perera, T. P. S. et al. Discovery and Pharmacological Characterization of JNJ-42756493 (Erdafitinib), a Functionally Selective Small-Molecule FGFR Family Inhibitor. Mol. Cancer Ther. 16, 1010–1020 (2017).

15. Hartley, R. S., Lewellyn, A. L. & Maller, J. L. MAP kinase is activated during mesoderm induction in Xenopus laevis. Dev. Biol. 163, 521–4 (1994).

16. Graves, L. M., Northrop, J. L., Potts, B. C., Krebs, E. G. & Kimelman, D. Fibroblast growth factor, but not activin, is a potent activator of mitogen-activated protein kinase in Xenopus explants. Proc. Natl. Acad. Sci. U. S. A. 91, 1662–6 (1994).

17. Lavoie, J. N., L’Allemain, G., Brunet, A., Müller, R. & Pouysségur, J. Cyclin D1 Expression Is Regulated Positively by the p42/p44 and Negatively by the p38/HOG Pathway. Journal of Biological Chemistry 271, 20608–20616 (1996).

18. Williams, O. M. et al. A Single Cell Atlas of the Newt Iris During Lens Regeneration. Preprint at 10.64898/2025.12.06.692619 (2025).

19. Muhammad, K. A., Nur, A. A., Nurul, H. S., Narazah, M. Y. & Siti, R. A. R. Dual-specificity phosphatase 6 (DUSP6): a review of its molecular characteristics and clinical relevance in cancer. Cancer Biol. Med. 15, 14 (2018).

20. Finch, P. W. et al. Purification and molecular cloning of a secreted, Frizzled-related antagonist of Wnt action. Proceedings of the National Academy of Sciences 94, 6770–6775 (1997).

21. Huber, O. et al. Nuclear localization of beta-catenin by interaction with transcription factor LEF-1. Mech. Dev. 59, 3–10 (1996).

22. Molenaar, M. et al. XTcf-3 transcription factor mediates beta-catenin-induced axis formation in Xenopus embryos. Cell 86, 391–9 (1996).

23. Behrens, J. et al. Functional interaction of beta-catenin with the transcription factor LEF-1. Nature 382, 638–42 (1996).

24. Kang, B. W. et al. Phosphodiesterase 5 inhibitor mirodenafil ameliorates Alzheimer-like pathology and symptoms by multimodal actions. Alzheimers Res. Ther. 14, 92 (2022).

25. Tsissios, G. et al. Characterizing the lens regeneration process in Pleurodeles waltl. Differentiation 132, 15–23 (2023).

26. Reyer, R. W. DNA synthesis and the incorporation of labeled iris cells into the lens during lens regeneration in adult newts. Dev. Biol. 24, 533–558 (1971).

27. Xie, Y. et al. FGF/FGFR signaling in health and disease. Signal Transduct. Target. Ther. 5, 181 (2020).

28. Marega, M., Chen, C. & Bellusci, S. Cross-Talk Between Inflammation and Fibroblast Growth Factor 10 During Organogenesis and Pathogenesis: Lessons Learnt From the Lung and Other Organs. Front. Cell Dev. Biol. 9, 656883 (2021).

29. Sousounis, K. et al. Eya2 promotes cell cycle progression by regulating DNA damage response during vertebrate limb regeneration. Elife https://doi.org/10.7554/eLife.51217 (2020) doi:10.7554/eLife.51217.

30. Schindelin, J. et al. Fiji: an open-source platform for biological-image analysis. Nat. Methods 9, 676–682 (2012).

31. Stringer, C., Wang, T., Michaelos, M. & Pachitariu, M. Cellpose: a generalist algorithm for cellular segmentation. Nat. Methods 18, 100–106 (2021).

32. Bolger, A. M., Lohse, M. & Usadel, B. Trimmomatic: a flexible trimmer for Illumina sequence data. Bioinformatics 30, 2114–2120 (2014).

33. Bray, N. L., Pimentel, H., Melsted, P. & Pachter, L. Near-optimal probabilistic RNA-seq quantification. Nat. Biotechnol. 34, 525–527 (2016).

34. Pimentel, H., Bray, N. L., Puente, S., Melsted, P. & Pachter, L. Differential analysis of RNA-seq incorporating quantification uncertainty. Nat. Methods 14, 687–690 (2017).

35. Brown, T. et al. Chromosome-scale genome assembly reveals how repeat elements shape non-coding RNA landscapes active during newt limb regeneration. Cell genomics 5, 100761 (2025).

